# Large-scale manipulation of radial positioning does not affect most aspects of genome organization

**DOI:** 10.1101/2024.08.22.609170

**Authors:** Mor Guetta, Hagai Kariti, Liora Sherman, Lena Israitel, Moran Tel-Paz, Nili Avidan, Revital Shemer, Noam Kaplan

## Abstract

Although the spatial organization of the genome is closely linked to biological function, genome structure is highly stochastic. Within this heterogeneity, specific function-associated structural elements must be maintained, even when the nucleus is deformed due to physiological physical constraints. The radial positioning of genomic loci - their distance from the nuclear periphery - has long been considered an important feature of genome organization which is correlated with both structural elements and genomic activity.

In the current study, we developed an experimental system for manipulating the radial positioning of the genome, by expressing the sperm-specific protein Prm1 in somatic cells. By microscopy, we observe that initial Prm1 nuclear foci develop within 72 hours into a large Prm1 focus occupying most of the nuclear interior, while the entire genome is driven towards the nuclear periphery, resulting in a 3-5 fold reduction in the volume that the genome occupies. Noting that this system enables isolation of a pure population of cells with reorganized nuclei, we then used Hi-C to study the effects of this perturbation. Remarkably, we find that interaction patterns are largely robust to this major nuclear reorganization, with minor changes which mostly reflect a strengthening of heterochromatin self-interactions.

Our experimental system provides means for manipulating nuclear organization in a reproducible manner, potentially allowing to examine radial positioning decoupled from other features of genome organization. Highlighting the complementary nature of microscopic and genomic methods, our work further suggests a remarkable resilience of genome structure such that large-scale nuclear changes, including chromosome compression and changes in radial positioning, can occur without extensive alteration of functional genome organization.

## Introduction

The spatial organization of the genome within the nucleus is highly stochastic, as reflected by a high level of heterogeneity within cell populations^1,2^. Despite this stochasticity, the 3D genome is closely linked to nuclear molecular processes, including transcription^3,4^, replication^5–7^, damage repair^8,9^ and cell division^10–14^. Thus, although many aspects of chromatin organization are highly variable and essentially unstructured, some genomic structures, which are often associated with biological function, tend to be consistent across the cell population and are mediated by the binding of specific molecules. Therefore, genomic structures such as genomic compartments^15,16^ and topologically associating domains (TADs)^17–21^ - both of which are associated with biological function - are naturally defined probabilistically rather than by a singular structure ^22^. In physiological settings, cells must maintain these function-associated genomic structures while being subjected to physical forces and constraints which could potentially distort the nucleus and its organization.

One feature of genome organization which has been associated with function is the radial positioning of loci, i.e. the distance from the nuclear lamina^23–25^. Heterochromatin tends to be located at the nuclear periphery, while euchromatin is located more towards the center of the nucleus. Accordingly, genes tend to be less active when they are more peripheral. At the chromosome level, while the locations of chromosomes within the nucleus are highly variable between cells, chromosomes do have preferred radial positionings, with short gene-rich typically inhabiting the center of the nucleus. Thus, radial positioning is considered an important feature of genome organization which is largely correlated with genome compartmentalization and chromatin state.

Genome organization has traditionally been studied by microscopy, providing easily interpretable direct visual observation in both fixed and live samples. However, the association of observed structures with specific genomic loci is typically to handful of tagged loci. Resolution has also been limited, albeit steadily improving. Lately, there have been important advances in on both fronts, with several methods utilizing highly multiplexed FISH ^26–30^ to enable visualization of hundreds of loci at super resolution in fixed samples. Still, a high-resolution genome-wide of entire mammalian chromosomes is currently beyond reach.

Since the rise of experimental genomic techniques based on next generation sequencing, the study of genome organization has been boosted significantly. Many of these are based on the principles of Chromatin Conformation Capture (3C)^31^, in which chromatin loci that are nearby each other within the nuclear space are physically linked and ligated to create a library of chimeric sequences. Such libraries, produced in either a genome-wide (e.g. Hi-C^15,32^, Micro-C^33,34^) or targeted fashion (e.g. 4C^35^, Capture-Hi-C^36^, HiChIP/PLAC-Seq^37,38^, MChIP-C^39^), are then sequenced to retrieve information regarding spatial proximity of genomic loci in a high-throughput manner. 3C-based methods provide large-scale, high-resolution data, structural data which is directly linked with sequences and other genomic features, which complements microscopy approaches. Importantly, these methods do not directly visualize the genome structure or actual geometric distances, but rather measure interaction frequency. Furthermore, they are typically performed on bulk populations rather than single cells, thus providing only limited insight on cell-to-cell variability. While single-cell variants exist, these are still technically challenging and limited in resolution ^40–44^.

While generally in agreement with each other, microscopy and genomic methods, in some cases the perspectives provided by these can be quite different^45,46^. Indeed, a few studies have showed instances in which dramatic changes in nuclear organization have very little effect on bulk Hi-C interaction maps. A study by Falk et al^47^. showed that although the nuclear architecture of mouse rod photoreceptors and of lamin B receptor null thymocytes is inverted, such that euchromatin is located at the nuclear periphery and heterochromatin occupies the center of the nucleus, changes in the Hi-C interaction maps are minor. Another example was provided by Sanders et al.^48^, who used low-salt swelling to expand nuclei more than twofold, again showing little effect on Hi-C interactions. Finally, during the final stages of spermatogenesis, the genome is restructured and massively condensed in a process called protamination, in which most histones are replaced with short basic protamine proteins. Intriguingly, some Hi-C maps measured in mouse sperm are very similar to those of somatic cells and do not reflect the expected changes of protamination, although it is yet unclear whether this is due to technical issues associated with the genome’s hyper condensed state^49,50^.

In the current study, we developed a biological system for manipulating radial positioning in somatic cells by expressing the sperm-specific protein Prm1. Following this major nuclear reorganization, we characterize how this perturbation affects genome organization via microscopy and Hi-C.

## Results

### A system for investigating Prm1 expression in somatic cells

In order to induce a large-scale rearrangement of nuclear organization, we expressed the sperm-specific protein Prm1 in human somatic cells. While the stages of spermatogenesis – including protamination - are highly regulated, a previous study suggested that expression of human Prm1 expression in sheep fibroblasts may be sufficient to induce genome protamination ^51^. To study the effects of human Prm1 expression in human somatic cells, we first transfected HEK293 cells with histone H2B tagged with GFP (H2B-GFP), followed by an additional transient transfection of Protamine1 tagged with mCherry (Prm1-mCherry) 24 hours later. Finally, we fixed the cells and DAPI-stained DNA at 24, 48 and 72 hours post-Prm1 transfection, and imaged H2B-GFP, Prm1-mCherry and DAPI using confocal microscopy (**Figure 1a**).

**Figure 1.**
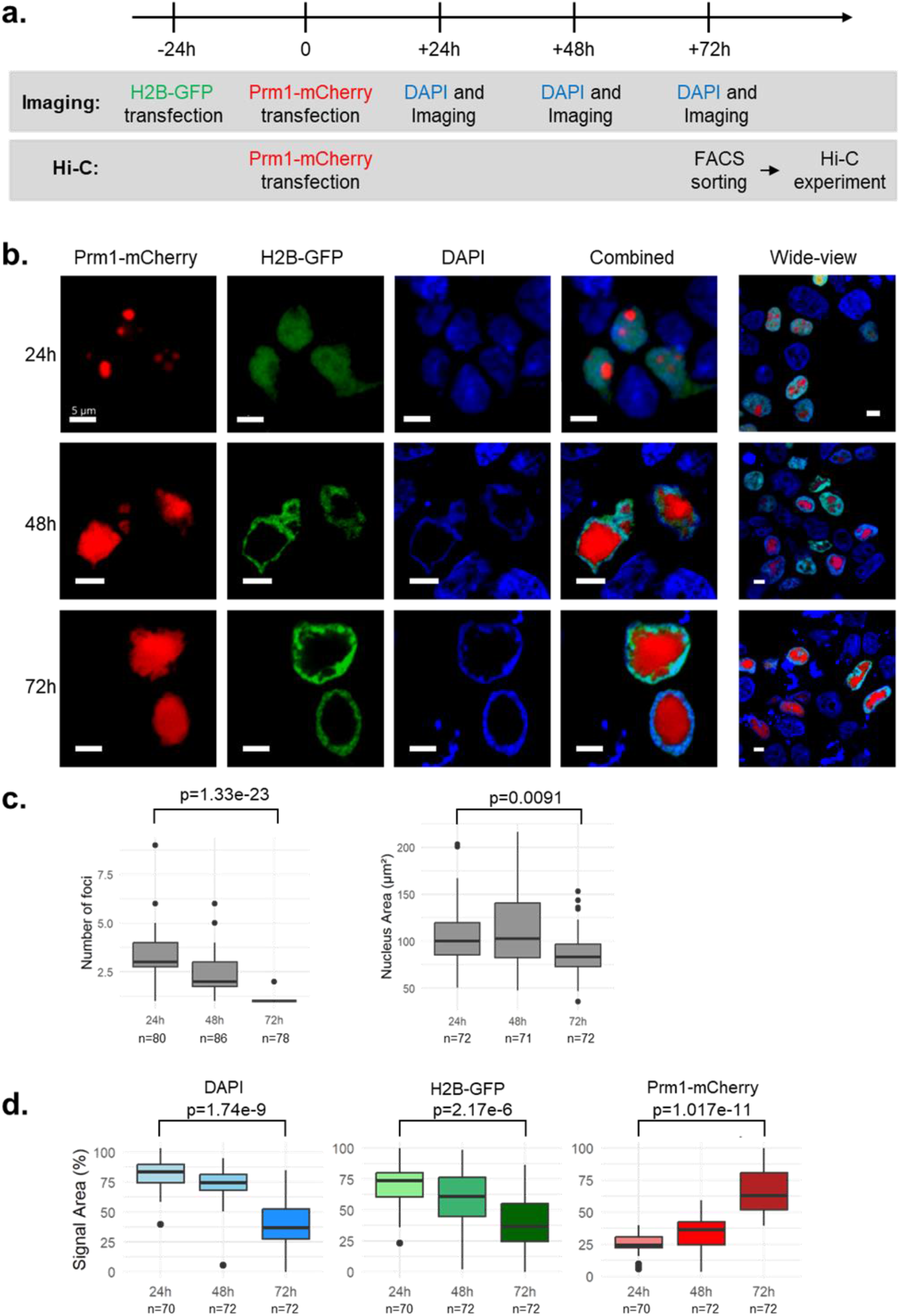
Prm1 expression in HEK293 cells. (a) experimental outline for microscopy and Hi-C. (b) Representative confocal projection images at 24, 48 and 72 hours post-Prm1 transfection (Red: Prm1-mCherry; Green: H2B-GFP; Blue: DAPI). (c) Quantification of number foci and nuclear area at 24, 48 and 72 hours post-Prm1 transfection. P-values are for Wilcoxon ranked sum test. (d) Quantification of relative nuclear area occupied by DAPI, H2B-GFP and Prm1-mCherry at 24, 48 and 72 hours post-Prm1 transfection. P-values are for Wilcoxon ranked sum test.

### Prm1 expression pushes the somatic genome towards the nuclear periphery

At the time of Prm1 transfection, nuclei showed standard nuclear organization in which DNA and H2B overlap and occupy most of the nuclear space, barring nucleoli. 24 hours post-Prm1 transfection, small Prm1-mCherry foci appeared scattered throughout the nuclei with minor overlap with H2B-GFP and DAPI (**Figure 1b, 1c**), consistent with the observations of Iuso et al. (2015). After 48 hours, the Prm1-mCherry appeared to form fewer but larger foci. possibly suggesting fusion of foci, with no co-localization of Prm1 with H2B-GFP or DAPI. Finally, after 72 hours, a single large Prm1-mCherry focus was observed at the center of most nuclei (**Figure 1b, 1c**). Notably, H2B-GFP and DAPI suggested that the chromatin had been pushed to the periphery of the nuclei. In contrast, in their study Iuso et al. (2015) observed co-localization of Prm1 and DNA signals with a simultaneous depletion of histone signal, suggesting DNA was protaminated. In our system, the absence of overlap between Prm1-mCherry and DAPI while H2B signal is maintained suggests that the chromatin has not been protaminated, but rather has been pushed to the periphery of the nuclei. To check that the absence of DAPI staining in Prm1-mCherry regions is not due to a high level of DNA condensation which prevents DAPI access to the DNA, we repeated Prm1-transfection after prestaining with live-cell DNA stain SPY-650 and observed similar results (**Figure S1**).

Over time, Prm1-mCherry occupied an increasingly larger area of the nuclei (on average 24.4%, 33.7% and 65.7% of the overall area after 24, 48 and 72 hours respectively) (**Figure 1d**). Accordingly, H2B-GFP and DAPI occupied a smaller area over time (69.7% and 81.6% after 24 hours, 60.1% and 72.1% after 48 hours, 37.3% and 38.6% after 72 hours). Thus, at 72 hours post Prm1 transfection, we observed a mean 2-3 fold reduction in the nuclear area covered by DAPI, which corresponds to a 3-5 fold reduction in volume (assuming a spherical nuclear shape). These results suggest that Prm1 expression in HEK293 cells leads to a large-scale nuclear reorganization, not by protamination of the genome but by pushing and compressing the genome at the nuclear periphery. Effectively, this experimental system allows consistent and reproducible manipulation of genome radial positioning.

### Isolation of a pure population of reorganized cells

Applying a bulk genomic method such as Hi-C to the population of transfected cells would be non-optimal due to the inclusion of cells in which the nuclear reorganization did not occur or did not complete. We postulated that in our system the strong Prm1-mCherry signal observed 72 hours after Prm1 transfection could be used to automatically isolate a pure population of cells in which the nuclear reorganization has occurred. To this end, we transfected HEK293 cells with Prm1-mCherry, and after DAPI staining we FACS-sorted the cells by both Prm1-mCherry and DAPI expression (**Figure 1A**). We observed a clear separation of the cells into two groups by Prm1-mCherry. Cells were sorted into three groups: Prm1^neg^, Prm1^pos^/DAPI^low^, and Prm1^pos^/DAPI^high^ (**Figure 2A**). Since for sorting cells were in suspension rather than grown on coverslips, we confirmed with microscopy that the sorted cells showed Prm1-induced genome reorganization similar to those we observed previously (**Figure 2B**). We also observed that the Prm1^pos^/DAPI^low^ and Prm1^pos^/DAPI^high^ looked similar in terms of nuclear organization. Thus, our system provides simple means for isolating a pure population of reorganized cells.

**Figure 2.**
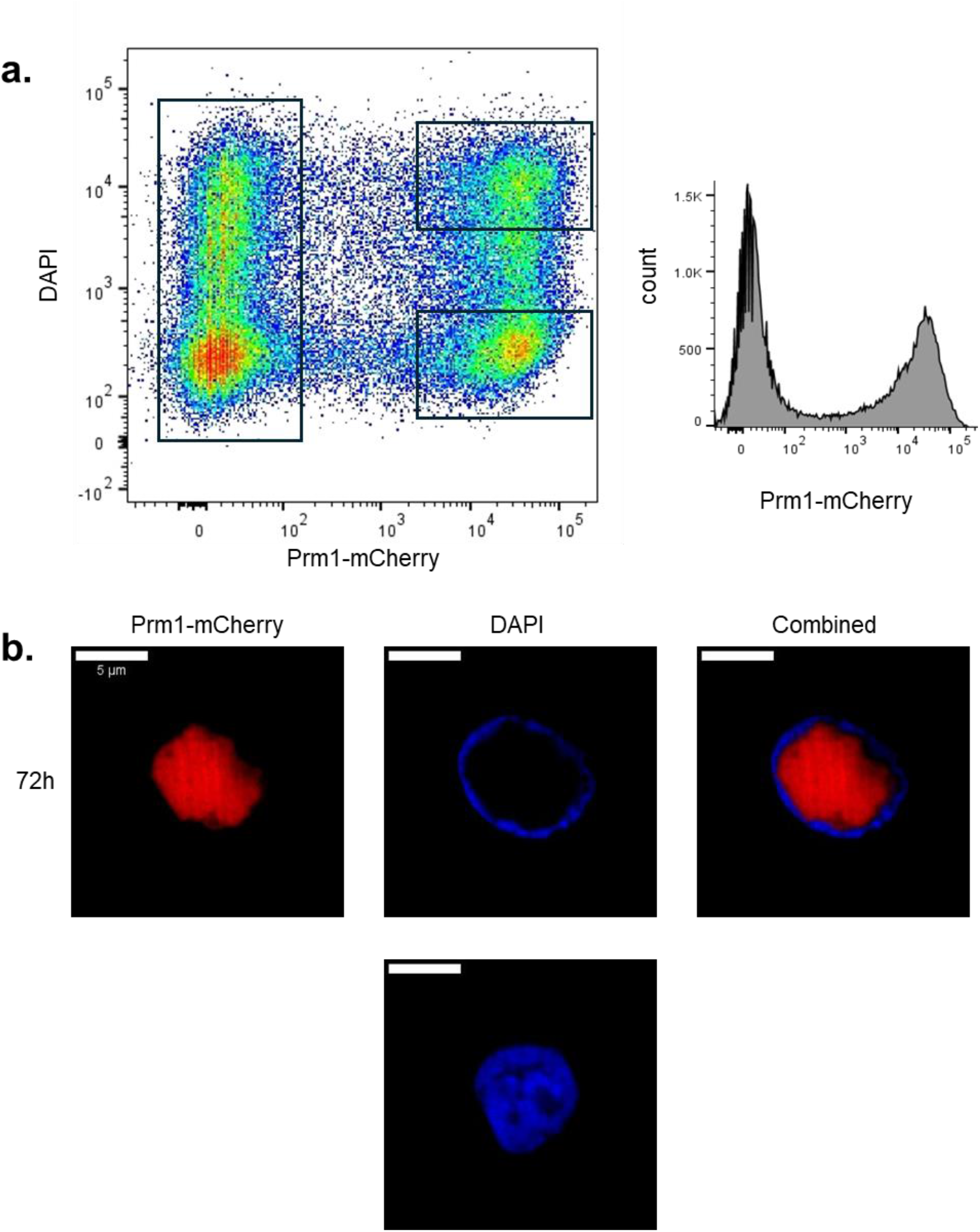
FACS separation by DAPI and Prm1-mCherry. (a) Left: FACS profile with rectangles indicating Prm1^neg^, Prm1^pos^/DAPI^low^ and Prm1^pos^/DAPI^high^ populations. Right: Histogram of Prm1-mCherry signal. (b) Confocal image of sorted Prm1^pos^ (top) and Prm1^neg^ (bottom) cells.

### Hi-C interaction patterns are globally maintained upon Prm1 expression

We next asked how the large-scale change in radial positioning induced by Prm1 expression affects 3D genome organization as reflected by interaction frequency patterns. We thus performed Hi-C on the sorted cell populations, sequencing a total of reads. All samples passed standard controls and quality metrics, and were mapped and processed to produce interaction maps. Since Prm1^pos^/DAPI^low^ and Prm1^pos^/DAPI^high^ interaction maps were highly similar, consistently with the microscopy results, we pooled these samples together and only considered Prm1^neg^ vs Prm1^pos^. To study the effect of Prm1 expression on 3D genome organization, we compared the interaction maps of Prm1^neg^ vs Prm1^pos^.

As an initial global comparison, we compared chromosome-level interactions between the samples. Previous analysis of interactions at this scale suggested that chromosome-chromosome interactions are associated with the radial positioning of chromosomes, while our own published model found that chromosome-level interactions are largely explainable by the composition of chromatin states on each chromosome^22^. Remarkably, we observe very little change in chromosome-level interactions upon Prm1 expression (**Figure 3**).

**Figure 3.**
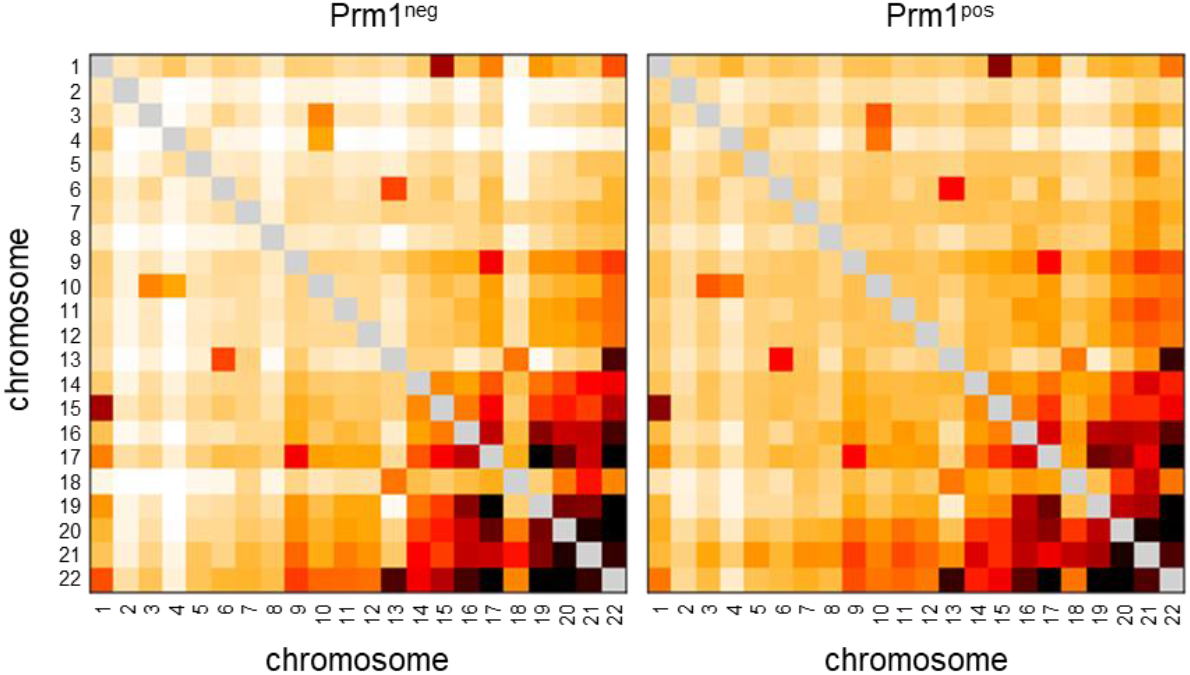
Chromosome-level interactions. Pairwise chromosome-chromosome average interaction frequencies for Prm1^neg^ and Prm1^pos^ populations. Intrachromosomal interactions are greyed out due to scale.

### Changes in distance-dependent interaction upon Prm1 expression

We next examined pairwise interaction frequency as a function of genomic distance. While ignoring many small-scale interaction patterns, analysis of distance-dependent interaction has proved to be useful in gaining general insight into fundamental polymer characteristics and distinguish between mechanistic models of chromosome folding ^10,11,52^. Plotting average interaction frequency versus genomic distance, we observe that Prm1^neg^ and Prm1^pos^ are overall similar, matching the previously observed approximately power law scaling of interaction frequencies (**Figure 4a**). However, we noticed small differences in the interaction profiles: at distances of up to a Mb (i.e. TAD scale), Prm1^pos^ shows fewer interactions than Prm1^neg^, while at larger distances (i.e. genomic compartment scale) Prm1^pos^ shows more interactions than Prm1^neg^ (**Figure 4a, 4b**). Due to the small magnitude of the differences, we compared at each genomic distance the average interaction frequency in each of the chromosomes of Prm1^neg^ versus each chromosome in Prm1^pos^. We find that these differences, although small, are statistically significant (paired t-test). Overall, these results suggest a slight increase in long-range interactions upon Prm1 expression, in line with the relative compression which the chromosomes undergo.

**Figure 4.**
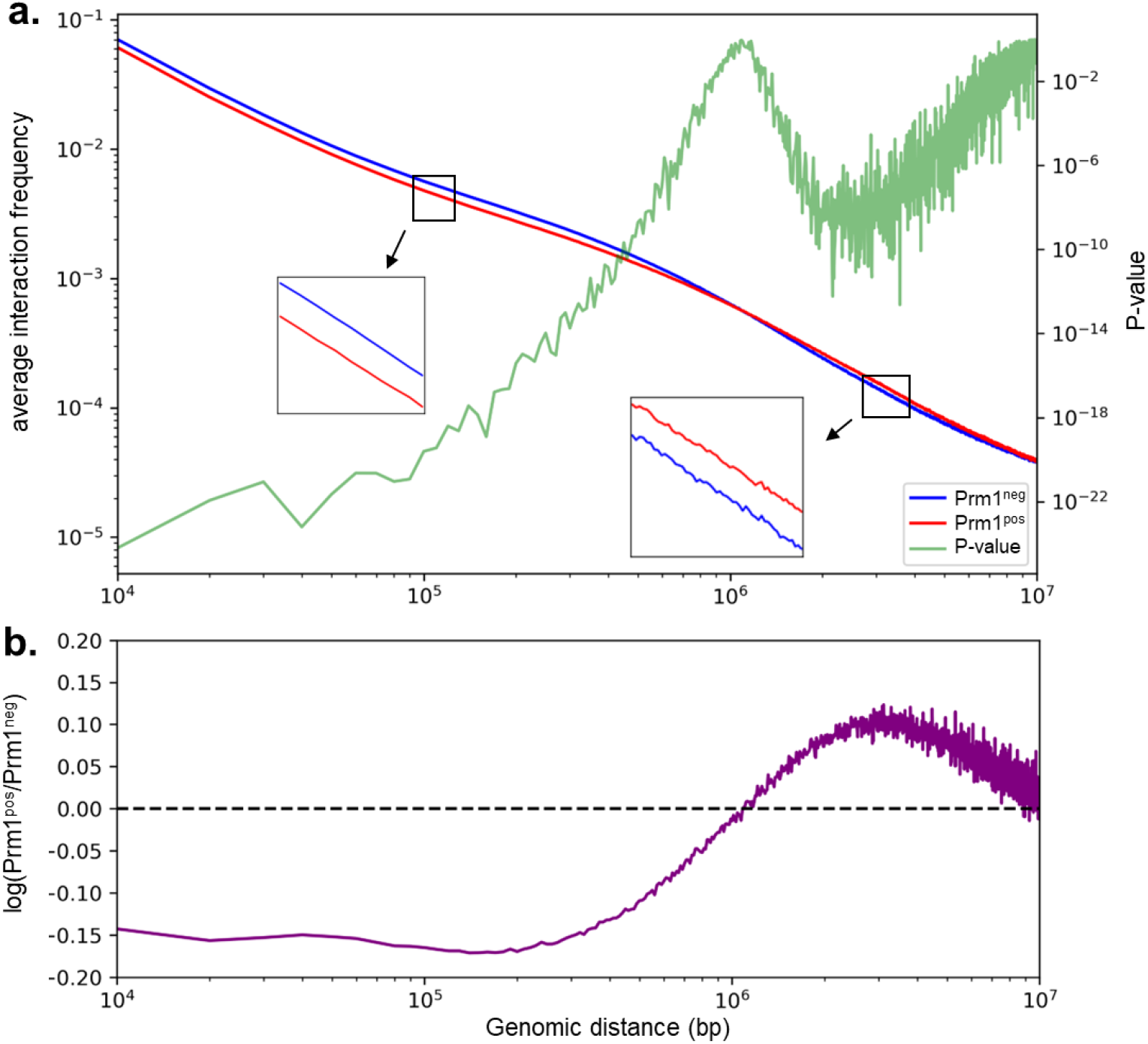
Distance-dependent interaction analysis. (a) Average interaction frequency as a function of genomic distance for Prm1^neg^ (blue) and Prm1^pos^ (red). Paired t-test p-value between the chromosomes of each sample at each genomic distance is shown in green. (b) log of the ratio between Prm1^pos^ and Prm1^neg^ average interaction frequencies at each genomic distance.

### Genomic compartmentalization is largely independent of radial positioning

Given the known association between radial positioning and genomic features including genomic compartments, we next asked whether Prm1 expression leads to changes in genome compartmentalization. To this end, we used deGeco, a probabilistic state model we have developed previously ^22^, to infer the state probability of each genomic locus, as well as the state-state affinities (note that the term affinity here is not used to imply actual physical force). We used a two-state model, which generally corresponds to the two states underlying standard A and B compartments (A: high probabilities; B: low probabilities). Remarkably, upon visual examination of the interaction maps, genomic compartment patterns looked highly similar in Prm1^neg^ and Prm1^pos^, and this was supported by the high correlation of their inferred state probabilities (Pearson r=0.983) (**Figure 5a**). However, Prm1^pos^ state probabilities were globally shifted to slightly lower values than those of Prm1^neg^, indicating a slight genome-wide increase in the tendency of loci to be in the B state (**Figure 5c**).

**Figure 5.**
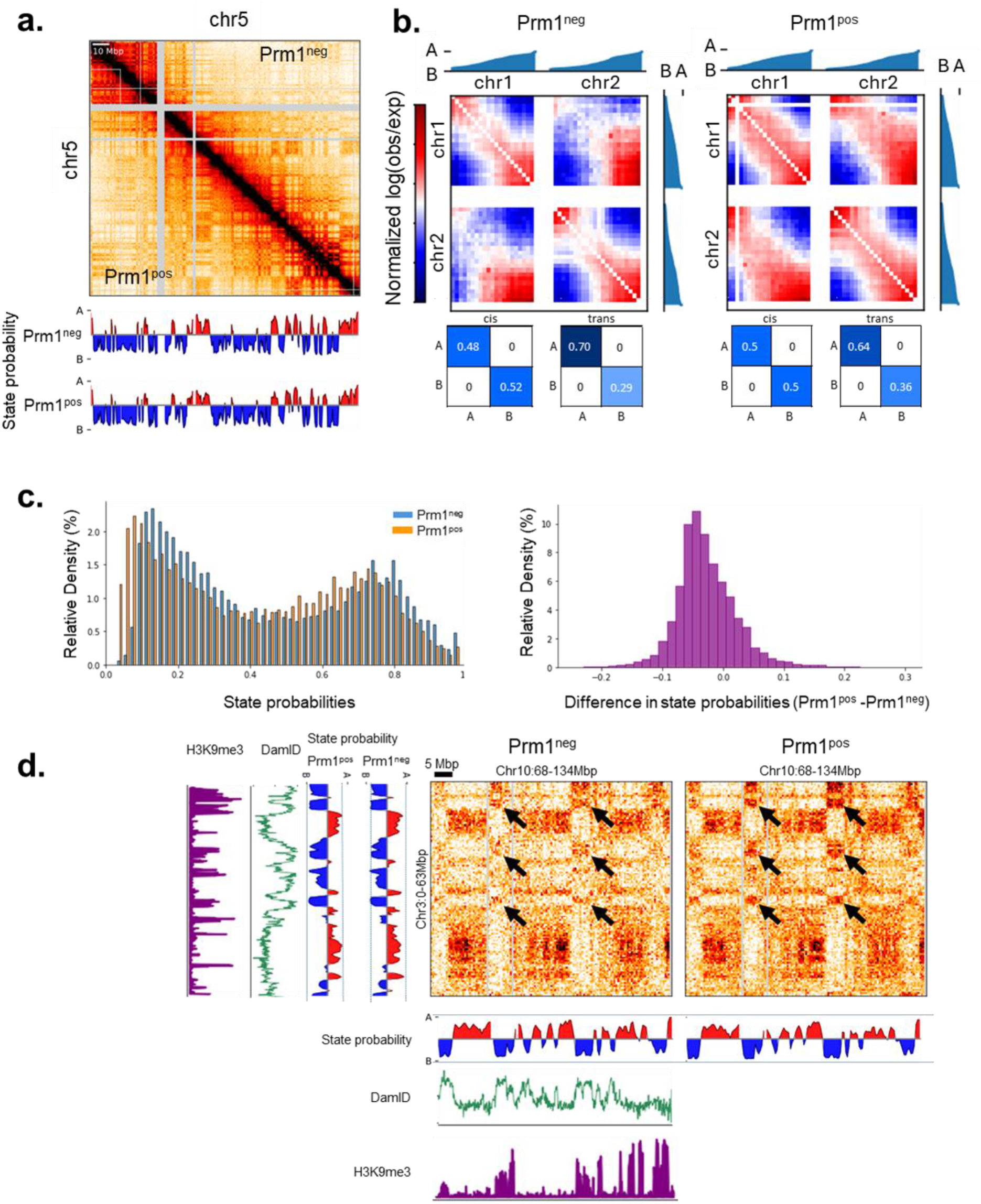
The effects of Prm1 expression on genome compartmentalization. (a) Interaction map of chromosome 5 for Prm1^pos^ and Prm1^neg^ alongside the inferred state probabilities. (b) Top: intrachromosomal (cis) and interchromosomal (trans) saddle plots, showing normalized log(observed/expected) interaction frequency after sorting loci by their state probability rather than their genomic location. Bottom: inferred state-state affinity cis and trans matrices for Prm1^pos^ and Prm1^neg^. (c) Histograms of state probabilities and difference in state probabilities. (d) Example of increased interchromosomal B-B interactions in Prm1^pos^, alongside genomic heterochromatin markers measured in wildtype HEK293.

As we previously found that inferred state-state affinities differ between intrachromosomal (cis) and interchromosomal (trans) interactions^22^, we asked whether this reorganization affects how states interact with each other (**Figure 5b**). We first verified that, as expected, in Prm1^neg^ the inferred affinities indicate that states A and B self-interact to a similar extent in cis (A-A 0.48 vs. B-B 0.52), while A-A is at least twice as strong than B-B in trans (A-A 0.70 vs. B-B 0.29). Upon Prm1 expression, we find that cis state affinities are maintained, in trans B-B affinity increases, while A-A affinity decreases (A-A 0.64 vs B-B 0.36). The strengthening of B-B interactions in trans was also apparent visually in interaction maps (**Figure 5d**), and matched characteristic heterochromatin features (lamin B1 DamID ^53^ and H3K9me3 ChIP-Seq ^54^). Thus, in spite of the entire genome being located at the nuclear periphery, the canonical self-interaction of A and B regions is not lost upon Prm1 expression, and even seems more pronounced in trans. In summary, our results suggest that the states underlying genome compartmentalization are generally resilient to the changes in radial positioning which are induced by Prm1 expression, with modest global changes in heterochromatin interaction levels.

### TAD structure is robust to Prm1 expression

Finally, we examined the effects of Prm1 expression at the scale of TADs. As a proxy of TAD signal, we calculated the insulation score ^55,56^, which is roughly the average amount of interactions across a genomic locus up to a defined genomic distance, for each locus. Following the previous observations at larger scales, we find that insulation is highly similar between Prm1^pos^ and Prm1^neg^ (Pearson r=0.939) (**Figure 6a**). While small differences in insulation score can be found, visual inspection of such regions shows they reflect more the limitations of the insulation score rather than meaningful distances in TAD structure (**Figure 6b**). In contrast to larger scale interaction patterns in which we found small yet consistent changes upon Prm1 expression, on the TAD scale we do not find such notable changes.

**Figure 6.**
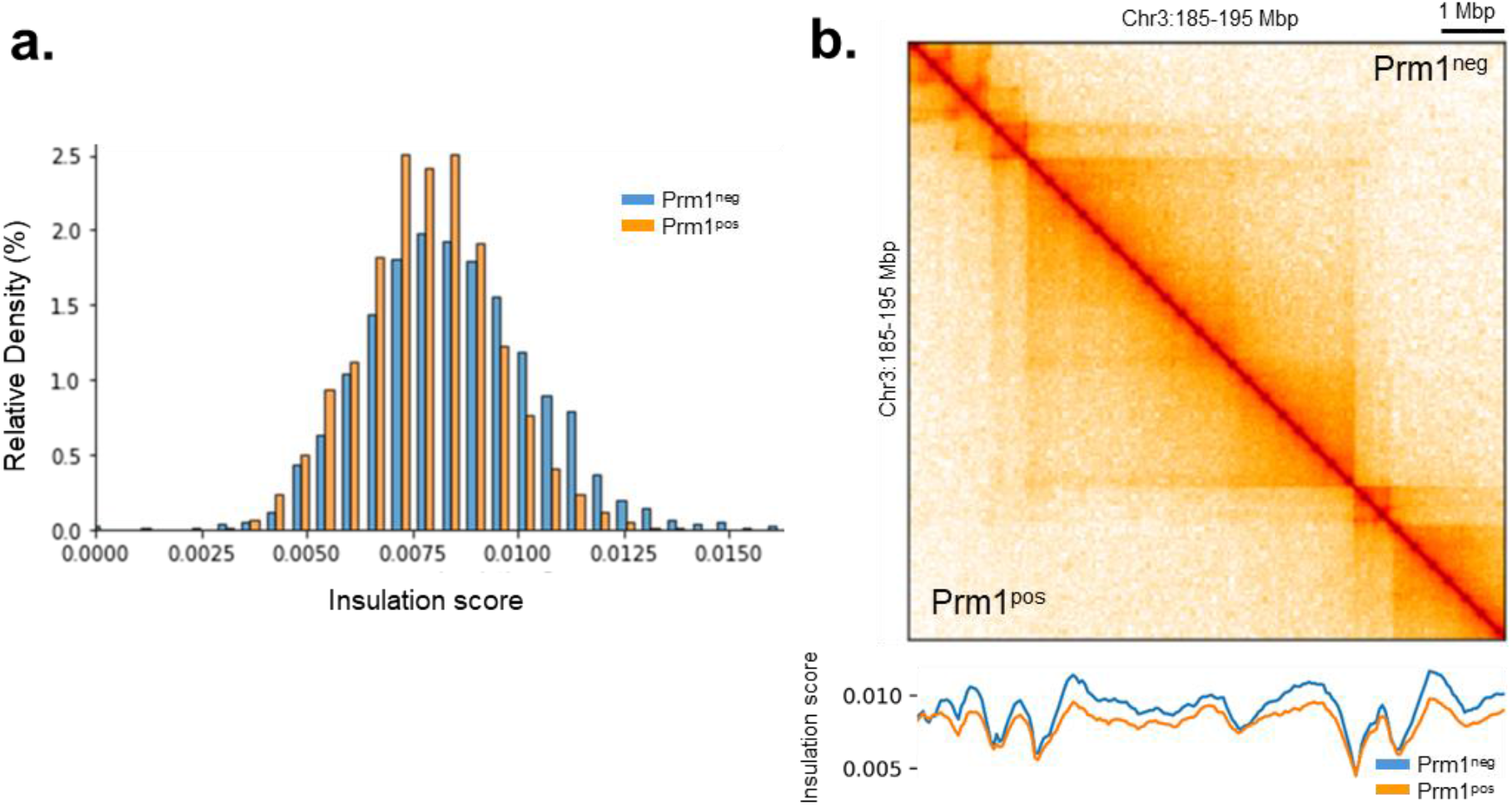
Effects of Prm1 expression on TAD structure. (a) Histogram of insulation scores for Prm1^pos^ and Prm1^neg^. (b) Example of minor changes in insulation score on TAD structure in Prm1^pos^ and Prm1^neg^ cells.

## Discussion

In this work, we developed a system which drives the genome to the nuclear periphery, thus changing its level of compression and its radial positioning. Our system allows high-throughput isolation of cells in which this reorganization has occurred, facilitating the application of follow up bulk assays. We use this system to study the connection between radial positioning and interaction frequencies. While the geometric positions and distances within the nucleus certainly must change under this perturbation, we surprisingly find that most of the structural features captured by Hi-C interaction frequency maps change very little. This apparent decoupling between radial positioning and genome organization, which is also reflected by the different perspectives provided by microscopy and genomic methods, suggests that the even upon deformation or compression, chromosomes are able to maintain an organization in which active and inactive regions self-interact without considerable mixing. While we do find a modest increase in heterochromatin interactions, likely due to proximity to the nuclear lamina, we do not observe any significant loss of the distinction between A and B compartments, as one might naively expect given that entire genome is located near the nuclear periphery. This does not necessarily mean that radial positioning in itself is unimportant, for example since factors entering the nucleus would reach regions in the nuclear periphery first. Indeed, this is the basis for GPSeq ^57^, a recent genomic method for estimating the radial positioning of genomic loci by break tagging following brief endonuclease digestions, which our work now shows can potentially measure a structural aspect of the genome which is not reflected in Hi-C interaction maps.

While the general checkered pattern of genomic compartments was maintained upon Prm1 expression, we were able to identify specific changes in compartmentalization. The changes we detected highlight the importance of the computational approaches used to analyze this type of data. We used deGeco, an explicit probabilistic model we developed previously^22^, which assumes that each locus in the genome belongs to one of *S* states (here we used *S*=2), and that pairs of loci interact according to their respective state probabilities while considering the intrinsic tendency of each pair of states to interact (which we refer to as state-state affinity). The model allows to infer these parameters, and they have a direct biological interpretation. Although the genomic profile of inferred state probabilities is typically quite similar to the first eigenvector produced by the standard PCA approach used for identifying genomic compartments, our model detects a global reduction in the state probabilities up on Prm1 expression, a phenomenon which would be difficult to identify using the eigenvector approach. In addition, the model allows us to reliably infer and detect changes in state-state affinities as we indeed observe in this case.

Our work is not the first instance of a large genome reorganization observed in microscopy which is barely noticeable in interaction maps^47,48^. Intriguingly, the changes we find in the profile on interaction frequency by genomic distance closely resemble the minor changes observed by Sanders et al.^48^ upon nuclear expansion. Is this a coincidence? More generally, it would be interesting to identify the common principles of these cases, in order to better understand what class of deformations and perturbations the genome can sustain without drastically changing its functional compartmentalization and TAD structure.

## Methods

### Plasmid Design and Preparation

Plasmid construction was performed using the NEBuilder HiFi DNA Assembly Cloning Kit following the manufacturer’s protocol. Sequences for the coding regions of human Prm1 coding region and mCherry, or human H2B and GFP, were synthesized and introduced into the SP-dCas9-VPR plasmid (Addgene Plasmid #63798, kindly provided by Izhak Kehat) after removing the CRISPR-associated sequences. Two plasmids were generated: pcDNA-hPrm1-mCherry, and pcDNA-hH2B-GFP. Both expressing the target gene under CMV promoter. The plasmids were sequenced for quality assessment and transformed into NEB® 5-alpha Competent Escherichia coli according to the NEBuilder protocol.

### Cell Culture

HEK293 cells (kindly provided by Raz Palty) were cultured under standard conditions. Cells were maintained in Dulbecco’s Modified Eagle Medium (DMEM) supplemented with 10% fetal bovine serum (FBS) and 1% penicillin-streptomycin. Cultures were kept at 37°C in a humidified atmosphere containing 5% CO_2_. Cells were passaged every 2-3 days to maintain exponential growth. Cell density was monitored to ensure optimal confluency of 70-80% before transfection experiments.

### Transfection

For microscopy analysis: Cells were plated in 24 wells, on coverslips coated with poly-L-lysine 24 hours before transfection. Transfections were performed using PolyJet™ In Vitro DNA Transfection Reagent, according to the manufacturer’s protocol. Initially, HEK293 cells were transfected with H2B-GFP plasmid. After 24 hours, the cells were washed with DMEM to remove excess plasmid, and a subsequent transfection with Prm1-mCherry plasmid was performed. Transfected cells were incubated at 37°C in a humidified atmosphere containing 5% CO_2_

For Hi-C experiments: HEK293 cells were plated 24 hours before transfection in 10 mm plates. They were then transfected with Prm1-mCherry plasmid and incubated for 72 hours before proceeding with the Hi-C sample preparation.

### Imaging

Imaging was performed at 24, 48, and 72 hours post-transfection. Cells were fixed with 4% formaldehyde for 15 minutes at room temperature, permeabilized with 0.1% Triton X-100 for 5 minutes, and blocked with glycine for 20 minutes. The coverslips were then stained with DAPI to visualize the nuclei, sealed with ProLong™ Gold Antifade reagent, and allowed to dry for 24 hours. For live-cell chromatin staining with SPY650-SiR-DNA, the cells were stained prior to transfection and imaged similarly post-transfection. Imaging was conducted using an LSM710 confocal microscope (Zeiss) or an IXplore Spinning disc confocal microscope (Olympus) at x40 and x60 magnification. Snapshots and Z-stack images were acquired to capture the entire nuclear volume. Each imaging experiment was repeated at least three times to ensure reproducibility.

### FACS Sorting

FACS sorting was conducted on cells transfected only with Prm1-mCherry, 72 hours post-transfection. Cells were suspended and fixed with 2% formaldehyde following the Hi-C protocol. After DAPI staining, cells were sorted into three populations using a FACS Aria™ Cell Sorter: cells with low mCherry (as negative control), cells with high mCherry and low DAPI signal, and cells with high mCherry and high DAPI signal. Sorted cells were then used for subsequent Hi-C experiments.

### Hi-C

Hi-C libraries were prepare using an optimized Hi-C protocol. Briefly, cells were crosslinked with 2% formaldehyde for 10 minutes at room temperature and quenched with glycine for 5 minutes. Cells were lysed in cold lysis buffer for 15 minutes on ice, followed by the addition of SDS and incubation at 65°C, then on ice, and Triton-X 100 and NEBbuffer were added, followed by incubation at 37°C. The nuclei were digested for 1.5 hours with DpnII restriction enzyme. End repair was performed with biotin-14-dATP, dNTPs, and Klenow enzyme, and incubated at 24°C for 1 hour. The biotin-labeled ends were ligated with T4 DNA ligase at 20°C for 1 hour. Crosslinking was reversed by adding proteinase K and incubating at 65°C for 1 hour. DNA was purified using phenol-chloroform and ethanol precipitation, then sheared by a Covaris M220 Focused Ultrasonicator. Biotin was removed from non-ligated DNA ends with T4 DNA polymerase. Biotinylated chimeric DNA strands were pulled down using Dynabeads™, and libraries were prepared using the NEBNext Ultra II DNA Library Prep. The libraries were checked with ClaI digestion and finally purified with AMpure XP magnetic beads before being sent for sequencing.

### Data Analysis

Microscopy images were processed and analyzed using ImageJ^58^, including measurement and quantification of signal area and size.

Hi-C libraries were processed using the distiller-nf pipeline (Version 0.3.4) to generate interaction maps in mcool format^59^. Briefly, paired-end reads were aligned to the human genome (hg38), uniquely mapping non-duplicate read pairs were retained, reads were binned into specified resolutions and interaction maps were filtered and balanced ^60^. Genomic compartments were analyzed with deGeco ^22^. All other Hi-C analysis was implemented in custom software. We used HiGlass ^61^ to visualize and explore the data.

HEK293 lamin B1 DamID was obtained from GEO GSE156150 ^53^. H3K9me3 HEK293 ChIP-seq data was obtained from ENCODE ENCFF190HHS ^54^. All datasets were binned into 100,000 bp intervals. Specifically, DamID data was originally binned, and ChIP-Seq data was aggregated by summing the values within each bin.

### Data Availability

Raw sequencing data and processed interaction maps (mcool) are deposited in GEO under GSEXXXXXX.

## Acknowledgements

We thank Jubran Boulos, David Cohen, Job Dekker, Izhak Kehat, Raz Palty, Lidan Shi, Haguy Wolfenson, the Technion Biomedical Core Facility imaging and genomics teams, and past and present members of the Kaplan lab. This research was funded by an Azrieli Early Career Faculty fellowship (NK).

## Notes

### Competing Interest Statement

The authors have declared no competing interest.

