## Supplemental Figures for "Large-scale manipulation of radial positioning does not affect most aspects of genome organization"

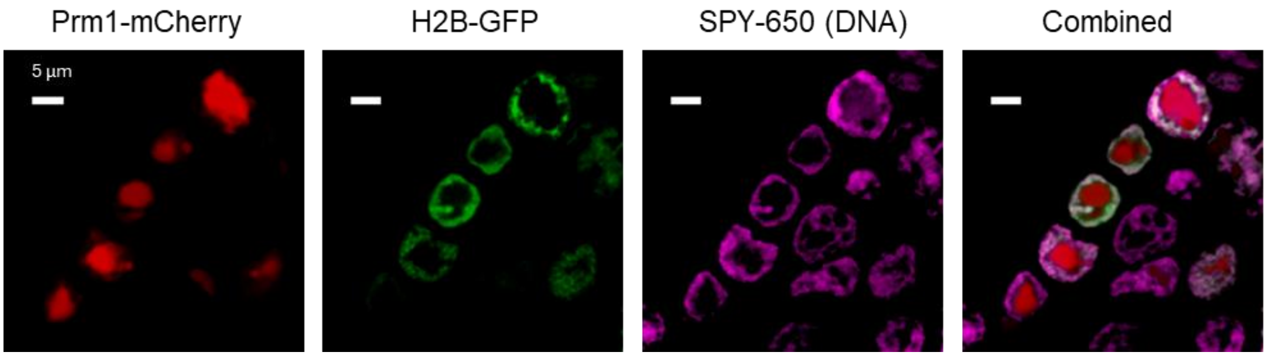


**Supplemental Figure S1. Genome reorganization with live-cell DNA stain.** Using the same protocol of transfection of HEK293 with H2B-GFP, after 24 hours we first stained DNA with SPY-650 and then proceeded to transfect with Prm1-mCherry. Example of nuclear reorganization at 72 hours after Prm1 transfection is shown.
